# SvAnna: efficient and accurate pathogenicity prediction for coding and regulatory structural variants in long-read genome sequencing

**DOI:** 10.1101/2021.07.14.452267

**Authors:** Daniel Danis, Julius O.B. Jacobsen, Parithi Balachandran, Qihui Zhu, Feyza Yilmaz, Justin Reese, Matthias Haimel, Gholson J. Lyon, Ingo Helbig, Christopher J Mungall, Christine Beck, Charles Lee, Damian Smedley, Peter N Robinson

**Author notes:** Shared first author (Daniel Danis, Julius O.B. Jacobsen).

## Abstract

Structural variants (SVs) are implicated in the etiology of Mendelian diseases but have been systematically underascertained owing to limitations of existing technology. Recent technological advances such as long-read sequencing (LRS) enable more comprehensive detection of SVs, but approaches for clinical prioritization of candidate SVs are needed. Existing computational approaches do not specifically target LRS data, thereby missing a substantial proportion of candidate SVs, and do not provide a unified computational model for assessing all types of SVs. Structural Variant Annotation and Analysis (SvAnna) assesses all classes of SV and their intersection with transcripts and regulatory sequences in the context of topologically associating domains, relating predicted effects on gene function with clinical phenotype data. We show with a collection of 182 published case reports with pathogenic SVs that SvAnna places over 90% of pathogenic SVs in the top ten ranks. The interpretable prioritizations provided by SvAnna will facilitate the widespread adoption of LRS in diagnostic genomics.

## INTRODUCTION

Structural variants (SVs) range from 50 base pairs (bp) to megabases in size and can be classified into a wide range of events including deletions, tandem and interspersed duplications, insertions, inversions, translocations, or complex combinations of these events.^1^ Limitations of existing sequencing technologies have hindered a full assessment of the role of SVs in human health and disease, but recent technological innovations are enabling more comprehensive detection of a broader range of SVs.^2^

The advent of short-read exome sequencing in 2010 ushered in a decade of novel discoveries in Mendelian genetics and led to the introduction of diagnostic exome and subsequently genome sequencing. Over 100 short read-based mappers and over 40 short-read variant callers have been introduced since 2010; while performance has been steadily increasing, SV calling from short reads is reported to have a recall of between 10 and 70% associated with high false-positive rates.^3,4^ Long-read sequencing (LRS), including both PacBio single-molecule real-time sequencing (SMRT) and Oxford Nanopore sequencing, produces longer reads that can be more accurately mapped to the reference genome even in regions that are inaccessible to short-read sequencing (SRS).^5^

LRS technology is rapidly evolving. A recent study with LRS in conjunction with additional methods such as single-cell template strand sequencing (Strand-seq) estimated that while 78% of SVs identified by SRS are concordant with LRS SV calls, only 30% of LRS SVs were observed in the short-read WGS callset; on average, 24,653 SVs were detected per genome by LRS.^6^ SRS captures the majority of SVs affecting coding sequence in genes with existing evidence for dominant-acting pathogenic mutations from OMIM, and the majority of SVs identified only by LRS are located in highly repetitive regions that have been previously inaccessible to human disease studies.^7^ Initial studies have appeared on LRS as a diagnostic tool for the diagnosis of Mendelian disease, with reports on the detection of large deletions, insertions, translocations, and tandem repeat expansions.^5,8–11^ LRS can be used to address cases in which SRS and sometimes cytogenetic or chromosomal microarray analysis has failed to identify an etiology, and therefore analysis may focus on intermediate size SVs (50 bp to 2 kb) difficult to detect with the other methods.^12^

In this work, we introduce SvAnna (see URLs), an integrated tool for the annotation and prioritization of SVs called in LRS data starting from variant call format (VCF) files produced by LRS SV callers such as pbsv, sniffles,^13^ and SVIM.^14^ SvAnna prioritizes variants in light of their overlap with structural elements of genes, promoters, and enhancers. On a curated set of 182 case reports of 188 SVs underlying Mendelian disease, SvAnna placed the correct variant within the top 10 ranks (out of 62,337-107,233 variants per VCF file) in 90.4% of cases.

## RESULTS

We developed SvAnna, a tool for phenotype-driven annotation and prioritization of SVs detected in LRS. SvAnna was designed to prioritize a broad range of SV classes such as deletions, duplications, inversions, copy number variants (CNVs), insertions, and translocations that affect one or more genes. SvAnna filters out common SVs and calculates a numeric priority score for the remaining rare SVs by integrating information about genes, promoters, and enhancers with phenotype matching to prioritize potential disease-causing variants. SvAnna outputs its results as a comprehensive tabular summary and as an HTML file intended for human consumption that visualizes each variant in the context of affected transcripts, enhancers, and repeats, providing information about the effects of the variant on transcripts, chromosomal locations, and Mendelian diseases associated with the affected genes.

### TADwise Assessment of Deleteriousness of Structural variation (TAD_SV_)

SvAnna assesses each variant in the context of the topological domain in which it is located. SvAnna first compares each variant to three sources of common SVs on the basis of reciprocal overlap. In addition to Database of Genomic Variants (DGV), gnomAD-SV, and dbSNP, which are largely based on data from SRS, SvAnna includes a recent dataset of SVs called by LRS^6^ (HGSVC). Common variants are removed according to user-defined frequency and overlap constraints (Methods).

SvAnna uses a set of evolutionary constrained, ubiquitous topologically associating domains^15^ (TADs) to determine the genomic neighborhood for assessing potential deleterious effects of SVs, an approach we term TADwise assessment of deleteriousness of structural variation (TAD_SV_). For each SV, SvAnna determines the extent of overlap with genomic elements, including enhancers, promoters, and transcripts. For each transcript, it determines which exon or exons are affected and whether the transcriptional start site of the coding sequence is disrupted. SvAnna then assembles a computational model of the TAD context of the variant. If a variant disrupts a TAD boundary, SvAnna defines a neo-TAD based on the non-disrupted TAD boundaries that flank the SV.

For each class of variant, SvAnna defines rules to assess the functional impact score *δ(G)* for a set of genes *G* located in the TAD (Figure 1A). At the same time, a phenotypic relevance score *Φ(Q,D)* is calculated based on the similarity of patient phenotypes *Q* encoded using Human Phenotype Ontology^16,17^ (HPO) terms and the ~8000 computational disease models *D* of the HPO project (Figure 1B). The candidates are ranked based on a TAD_SV_ score that is calculated as a function of the *δ(G)* and *Φ(Q,D)* scores. The following sections explain the approach to specific classes of SV.

**Figure 1.**
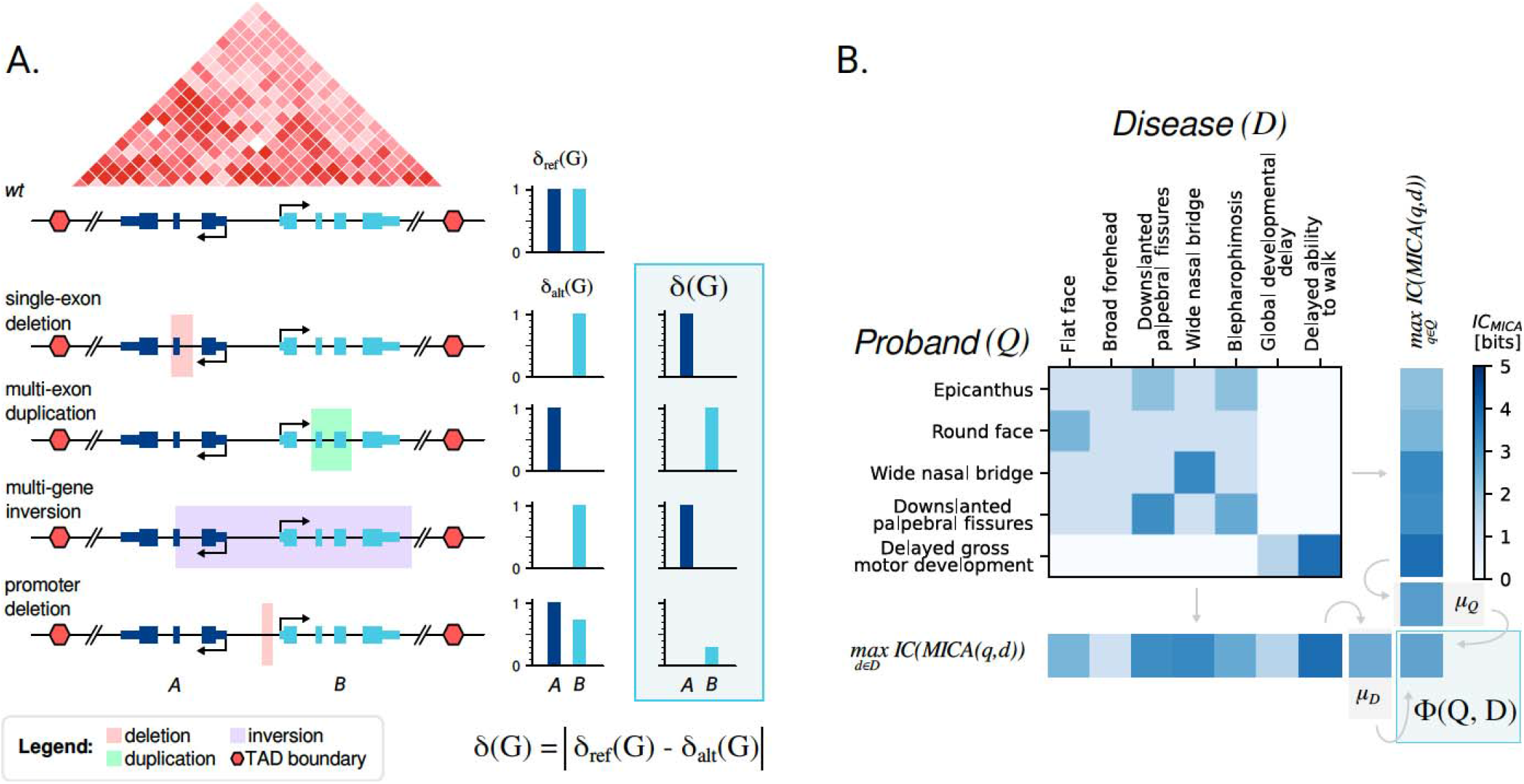
Overview of SvAnna algorithm. **A. Functional impact score *δ(G)***. The score compares reference and alternate sequences. For the reference, each gene *g* in the TAD is assigned a *δ_ref_(g)* score of 1. A deletion of a coding exon or an entire gene *g* is assigned a *δ_alt_(g)* score of 0. A duplication is assigned a score of 0 if it disrupts the coding sequence of a gene, and 2 if it completely encompasses the gene. A coding sequence that is disrupted by an inversion is assigned a score of 0, but a gene that is completely contained within an inversion is assigned a score of 1. A deletion in a promoter sequence is assigned a score based on the length of the deletion. The *δ(G)* scores for a variant are calculated as the absolute difference between wildtype *δ_ref_(G)* and variant *δ_alt_(G)* scores. **B. Phenotype similarity score *Φ(Q,D).*** SvAnna calculates the phenotypic similarity for a set of HPO terms *Q* representing the patient’s phenotypic features and HPO terms *D* for a disease. SvAnna computes the information content (IC) of the most informative common ancestor (MICA) for all term pairs *q,d* for *q ∈ Q* and *d ∈ D*. The mean ICs *μ_Q_* and *μ_D_* are calculated for Q and D, and the final similarity score *Φ* is calculated as the mean of *μ_D_* and *μ_Q_.* The *δ(G)* and the *Φ(Q,D)* scores are combined to obtain the final TAD_sv_ score (Methods).

To demonstrate SvAnna, we curated 182 published case reports with 188 distinct SVs associated with Mendelian disease. Phenotypic data in the reports was captured as HPO terms and recorded together with genotypic data using the Global Alliance for Genomics and Health (GA4GH) Phenopacket format.^18^ For each case, the causal variant was added to an in-house whole-genome sequencing VCF file (Methods) called by the PacBio structural variant caller pbsv (see URLs), and the phenotypic descriptions were encoded using HPO terms (Table 1 and URLs). The analysis was repeated using 10 different in-house VCF files and the median rank was recorded.

**Table 1:** Summary of curated collection of deleterious SVs. We curated a collection of 188 published deleterious SVs based on 182 cases published in 146 clinical case reports. We considered five classes of SVs commonly present in LRS variant calling results: deletions, duplications, inversions, insertions, and translocations. We further classified the SVs into five functional categories based on the number of affected genes and the relative location of the SV region with respect to transcripts of genes. The case reports are available for download (URLs).

|  | Deletions | Duplications | Inversions | Insertions | Translocations | Total |
| --- | --- | --- | --- | --- | --- | --- |
| Multiple genes | 7 | 2 | 4 | N/A | 5 | 18 |
| Multiple exons | 37 | 19 | 5 | N/A | N/A | 61 |
| Single exon | 81 | 16 | 2 | 3 | N/A | 102 |
| Promoter | 5 | 0 | 0 | 0 | 0 | 5 |
| Intronic | 2 | 0 | 0 | 0 | N/A | 2 |
| Total | 132 | 37 | 11 | 3 | 5 | 188 |

### Deletions and duplications

To calculate the priority of a deletion of a genomic region, SvAnna determines the relative location of each overlapping gene with respect to the variant. Deletions can be located within a transcript region, completely contain a transcript, or partially overlap. SvAnna assigns complete transcript deletions a *δ(g)* score of 0. Deletions of single exons are assigned a score of 0 if they include coding sequence or a canonical splice site region. If a deletion is located in an intron it is assigned a score of 1 (not deleterious) except if it overlaps with an enhancer sequence (see below). If a deletion affects an untranslated region (UTR), it is assigned a score based on the length of the SV and the UTR, with lower (more deleterious) scores being assigned to SVs that are large compared to the UTR sequence. In some cases, the effect of an SV is different for different transcripts of a gene. SvAnna chooses the most deleterious (lowest) score for any transcript of a gene and uses that score to calculate the *δ(g)* score. If a deletion encompasses multiple genes, the *δ(g)* score is assigned in this way for each transcript of each gene.

For example, a ~6.9kb-long deletion that leads to in-frame loss of 48 amino-acid residues encoded by exon 2 of *NF1* (NM_000267.3)^19^ is considered to abolish the function of the gene *(δ(g)=0)* as the deletion removes the entire exon from the transcript. Together with the phenotype features of the proband consisting of *plexiform neurofibroma* (HP:0009732), *spinal neurofibromas* (HP:0009735), *tibial pseudarthrosis* (HP:0009736) and *multiple cafe-au-lait spots* (HP:0007565), the variant attains rank 1 with a final TAD_SV_ score of 117.4 (**Figure 2A**). SvAnna generates an HTML output of the variant and its position with respect to the affected transcripts, repetitive elements, and enhancers (see **Supplemental Fig. S1** for this example). The median rank of 81 cases with single exon deletions was 1.

**Figure 2.**
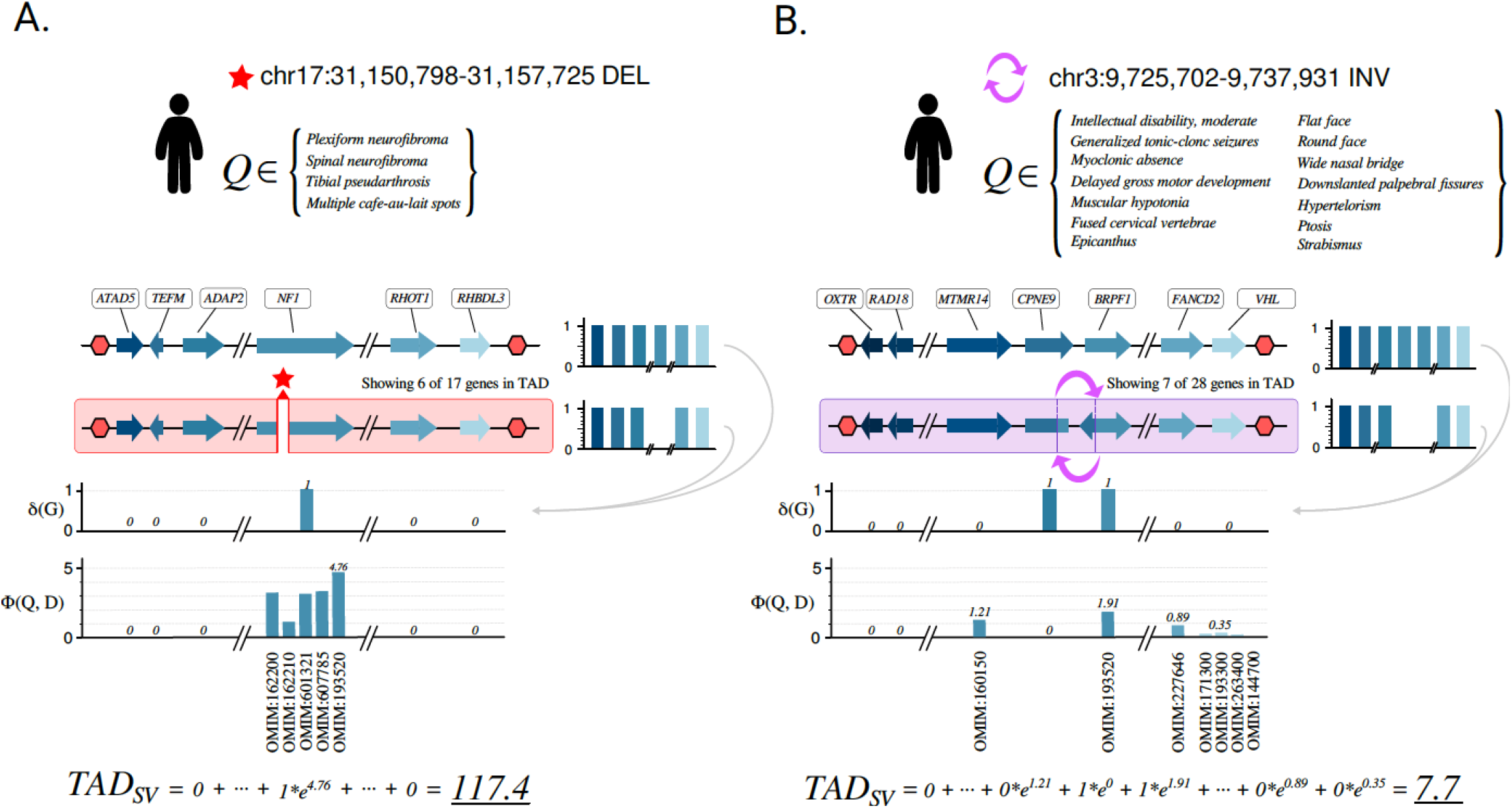
Prioritization of variants. **A.** A case of proband with a single-exon deletion in the *NF1* gene.^19^ SvAnna calculates *δ(G)* scores for all 17 genes *G* located in the TAD region. The deletion results in *δ(g)=1* for *NF1*. To calculate semantic similarity *Φ(Q,D)* for *NF1,* SvAnna evaluates five computational disease models associated with variants in *NF1*. In case of this proband, *Neurofibromatosis, Type I* (OMIM:193520) is the disease model that matches the proband’s clinical condition the best (Φ *(Q,D)* = 4.76). As *NF1* is the only affected gene in the TAD, *δ(g)* and *Φ(Q,D)* of *NF1* are the only determinants of the final TAD_SV_ score. **B.** A case of proband with an inversion involving 3’ end of *CPNE9* and 5’ end of *BRPF1*.^20^ SvAnna assigns *δ(g)* score of 1 to both *CPNE9* and *BRPF1* that are disrupted by the inversion. Unlike in the case of *NF1* variant, more genes are potentially phenotypically relevant to proband’s clinical description. The final TAD_SV_ integrates the scores of phenotypically relevant *BRPF1* (6.7) and disrupted, but phenotypically non-relevant *CPNE9* (1).

A deletion of three exons in *BRCA1* received a TAD_SV_ score of 7.76 and rank 1 (**Supplemental Fig. S2**). The median rank of multiple-exon deletions was 1. SvAnna uses a slightly different approach to prioritize multigene SVs. For instance, a deletion at chr2:109,923,337-110,405,062 (hg38) affects four genes *(MTLN, MALL, MTLN,* and *NPHP1).* SvAnna calculates the *δ(g)* score of each gene as 1, weighted by the phenotype score according to equation 5 (Methods). Only *NPHP1* is associated with a phenotypically relevant disease (Joubert syndrome 4) and its contribution to the final TAD_SV_ score is highest (**Supplemental Fig. S3**). The median rank of deletions affecting multiple genes was 1.

Duplications are handled in a similar way except that the gene *g* that is entirely spanned by a duplication is assigned *a* score of 2 and the *δ(g)* score is calculated as the absolute value of the difference between the wildtype (1) and the variant (2). Similar considerations about duplications that affect individual exons or an entire transcript pertain as with deletions. Tandem duplications that do not alter the primary linear sequence of a transcript (e.g., a duplication of the final exon of a transcript) are assigned a score of 1 (i.e., are assumed not to be deleterious). For example, a pathogenic duplication of 36 bp within one exon of the *PIBF1* gene was assigned a TAD_SV_ score of 2.36 (**Supplemental Fig. S4**).

### Inversions

An inversion is prioritized using a similar approach to that used for evaluating a deletion, with several differences. The *δ(g)* score for a genomic element spanned by an inversion is defined as 1 (since the primary sequence of the element is unchanged). If the sequence of a transcript or enhancer is interrupted by inversion breakends a *δ(g)* score of 0 is assigned. Additionally, inversions that affect one or multiple (but not all) exons of a transcript are assigned a score of 0. An inversion located completely within an intron is assigned a score of 1 (not deleterious).

For example, breakpoints of a 12kb copy neutral inversion identified in monozygotic twins suffering from intellectual disability disrupt genic regions of *BRPF1* and *CPNE9*.^20^ Since each breakpoint disrupts the gene sequence, the *δ(g)* score is set to *0* for both genes. Deleterious variants in *BRPF1* are associated with *Intellectual developmental disorder with dysmorphic facies and ptosis* (OMIM:617333). The final TAD_SV_ score of 7.7 aggregates scores of two disrupted genes: phenotypically relevant *BRPF1* (6.7) and disrupted, but phenotypically not relevant *CPNE9* (1.0) (**Fig. 2B, Supplemental Fig. S5**).

### 5’UTR and transcriptional start site variants

SvAnna has specific rules for prioritizing SVs in non-coding sequences. SvAnna assumes that variants affecting UTR regions, especially SVs, are less likely to have a functional impact on gene expression or translation. SvAnna calculates the *δ(g)* score for a UTR variant as a function of the variant length and UTR length (Methods). However, the variants that disrupt transcription start sites (TSS) are considered just as deleterious as variants that affect coding sequences. As an example, we evaluated *de novo* deletion of 1,571 bp affecting the first non-coding exon of *AMER1*.^21^ The deletion was processed as a loss of TSS, leading to a TAD_SV_ score of 8.99 (**Supplemental Fig. S6**).

### Promoter variants

SvAnna extends the prioritization rules to variants in gene promoters with a potential to change the gene expression. The promoter regions are assumed to encompass 2kb upstream of the TSS. To calculate the *δ(g)* score, SvAnna assigns SVs in promoter regions a score that is 40% of that of an SV in a coding sequence. Effectively, this means that only promoter variants in genes with a good phenotype match get high priority scores. This is a limitation of the approach that could be overcome as our ability to build computational models of promoter variants improves. If the variant affects a promoter and another genic region (e.g. TSS, UTR), the most deleterious *δ(g)* score for any of the regions is used to calculate the TAD_SV_ score. For example, a 13-bp deletion in the promoter of the von Willebrand factor *(VWF)* gene in a patient with type 1 von Willebrand disease^22^ was assigned a TAD_SV_ score of 25.6 (**Supplemental Fig. S7**).

### Translocations

SvAnna applies a series of rules to assess the pathogenicity of translocations. A translocation that disrupts the coding sequence of a gene *g* or separates the transcription start site of *g* from its coding sequence is assigned a *δ(g)* score of 0. The TAD_SV_ score is calculated based on the predicted effects of the translocation at both breakpoints. For example, a translocation that disrupts the coding sequence of *SLC6A1* in a case of myoclonic-atonic epilepsy was assigned a TAD_SV_ score of 4.63 (**Supplemental Fig. S8**).

### SvAnna achieves clinically relevant sensitivity

To demonstrate the practical utility of the SvAnna algorithm, we curated a collection of 182 case reports with 188 known disease-causing SVs associated with Mendelian diseases (see URLs). In addition to genomic coordinates and genotypes of the causal variants, we recorded NCBI Gene and OMIM identifiers for the causal gene and the associated disease, and we encoded the clinical features of the proband into Human Phenotype Ontology terms.^16,17^ We stored the cases in Phenopacket format. The curated SVs included deletions, duplications, inversions, insertions, and translocations affecting a differing number of genomic elements.

We are not aware of any other tool that specifically targets VCF data as produced by modern LRS SV callers. We ran SvAnna on 10 in house VCF files called with pbsv, sniffles,^13^ and SVIM,^14^ and were able to annotate ~99.9% of variants (i.e., identify nature and position of overlaps with transcripts). We are aware of only one other published tool for phenotype-driven variant prioritization: *AnnotSV*,^23^ a standalone command-line script that annotates SVs with functional, regulatory, and clinical information to filter out neutral variants and rank the candidate pathogenic variants, while the phenotype matching is delegated to a separate tool, Exomiser.^24^ *AnnotSV* is able to annotate only DEL and DUP calls and missed INS, BND, CNV, and INV calls that were processed by SvAnna. For instance, on a VCF file called by pbsv, AnnotSV missed 25,198 (39.9%) or 63,084 variants.

To assess the practical utility of SvAnna and to compare performance with *AnnotSV,* we developed a simulation strategy based on 182 curated case reports. We used 10 VCF (62,337-107,233 variants) generated by the LRS pipeline and added the causal variant(s) to simulate 10 runs per curated case, for a total of 1,820 data sets. Then, we prioritized the simulated variant dataset and calculated the median rank for the causal variant across 10 runs. Overall, SvAnna placed the causal variant on the top of all variants in 61.2% of cases, the causal variant was at rank 10 or better in 90.4% of cases, and 93.6% of variants were placed at a rank of 20 or better (**Fig. 3A, B**).

**Figure 3.**
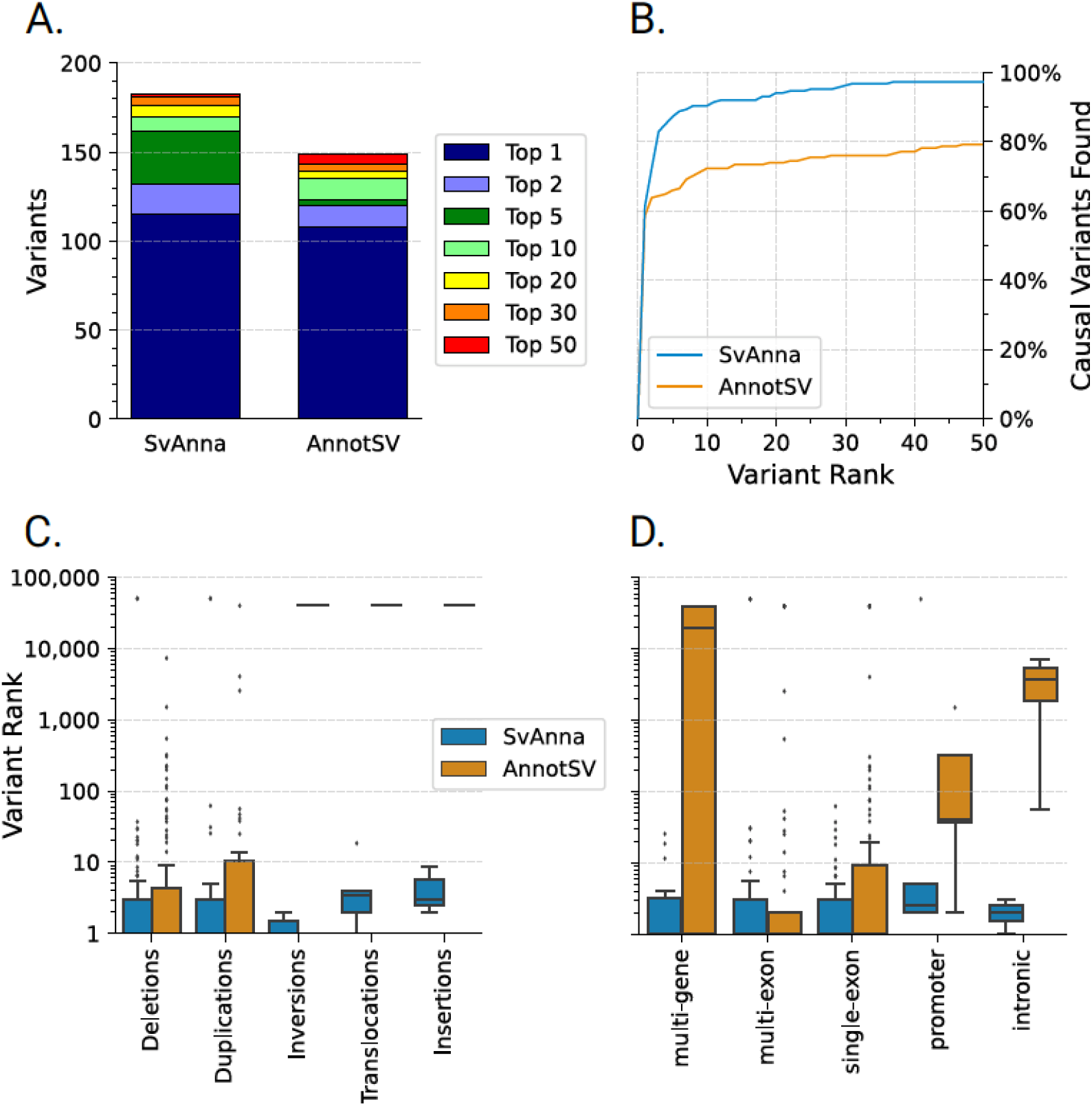
Comparison of prioritization performance of different methods for prioritization of SVs. **A.** Median ranks of 188 deleterious SVs obtained from simulated analysis runs. Top 5 means that the rank assigned by the tool was between 1 and 5, and so on. **B.** Plot showing the cumulative rank for prioritizations by SvAnna and AnnotSV. **C.** SvAnna assigns the best rankings to all 5 evaluated SV classes. **D.** SvAnna attains the best median ranks for SVs of all sizes, performing notably well in prioritization of variants involving multiple genes. In panels C and D, each box plot is defined so the center line is at the median variant rank, the box borders mark the 25th and 75th percentiles, and the whiskers stretch to denote 1.5 times the interquartile range.

We further evaluated the performance on different variant types. SvAnna showed consistent performance for all variant types from our benchmark set. SvAnna was confident in prioritization of deletions and duplications, assigning median variant rank of 1. The inversions and breakend variants had median rank 2 and 3.5 respectively. Insertions were the hardest to prioritize with a median rank of 5. SvAnna supports prioritization of all variant types (**Fig. 3C, D**).

### SvAnna software

SvAnna presents its results both as tabular and VCF files suitable for bioinformatics analysis as well as a visually appealing HTML report to support clinical interpretation. The HTML report is a single page with a tabular and graphical summary of the top 100 variants, with information about the variant (read counts, VCF id, position and length, and genotype if available), genes that overlap the variant and Mendelian diseases associated with the genes,^25^ and a list of all overlapping transcripts as well as the position and effect on the transcript. A graphical display is generated for each variant (such as those shown in **Supplemental Figs. S1-S8**) as a scalar vector graphics (SVG) file that is embedded directly in the HTML code. The SVG shows the SV and its position compared to that of overlapping transcripts, whose coding exons are shown in green and noncoding exons in yellow. If applicable, overlapping repeats^26^ and VISTA enhancers^27^ are shown as tracks beneath the variant (Screenshot in **Supplemental Fig. S9**). SvAnna is made freely available for academic use as a Java command-line application. The GitHub repository contains source code, a pre-built executable, and links to detailed instructions for use, as well as a VCF file with the eight examples presented here and a tutorial.

## Discussion

Our ability to analyze the role of SVs in Mendelian disease has lagged significantly behind that of single nucleotide variants and other small variants for a myriad of reasons including technical difficulties in calling SVs, the relative lack of functional data on the effects of SVs on gene regulation, the paucity of genome-wide association studies for SVs.^1^ The advent of LRS promises to greatly improve the detection of SVs in patient samples. PacBio LRS was shown to be about three time more sensitive than a SRS ensemble calling approach, with the improvement was predominantly derived from improved detection of repeat-associated SV classes, particularly of intermediate-sized SVs (50 bp to 2 kb), and insertions across the SV size spectrum.^28^ However, progress on many fronts will be required to fully realize the promise of LRS for genetic medicine, including continued technical improvements, cost reductions, better SV calling algorithms, and more comprehensive knowledge of the medical relevance of classes of SVs that were difficult or impossible to ascertain with previous technologies.

Compared to the large variety of approaches available for SRS, there are very few computational methods for assessing the relevance of SVs for rare disease.^2^ Numerous algorithms, databases, and tools have been developed to support the medical interpretation of diagnostic SRS. Although details vary from tool to tool, in general, variant pathogenicity is assessed on the basis of variant allele population frequencies, evolutionary conservation, and functional impact prediction for missense, splice, and regulatory variants.^29–31^ Disease genes can be prioritized based on functional and genomic data,^32^ or on the basis of phenotypic similarity of patient phenotype definitions with computational disease models of the HPO project.^16,33^

There is a need to extend these algorithms for LRS. SvAnna includes a number of innovations to this end. VCF files represent SVs in multiple ways including the default (sequence) representation and symbolic notation. Translocations are represented as breakend calls on two lines. SvAnna uses a harmonized computational model of variants to represent each category of variant, which enables it to apply a single prioritization approach to all categories of SV. We are aware of only one previously published tool for phenotype-driven prioritization of SVs, AnnotSV.^23^ In contrast to SvAnna, AnnotSV was primarily designed to analyze SV events identified in SRS and array-based experiments, and only supports the analysis of deletions and duplications. SvAnna demonstrated a substantially better overall performance than AnnotSV in the ranking of all classes of the causal SVs.

In our study, SvAnna prioritized over 90% of SVs in the first 10 ranks. The case reports were chosen from 182 publications, seven of which reported diagnostic results from LRS.^5,10,12,20,34–36^ Current published experience with LRS in a human genetic diagnostic setting is limited, and it is too early to assess the potential advantage of LRS over SRS in diagnostic settings. Given that LRS detects SVs in genomic regions that were difficult or impossible to characterize by SRS, the medical relevance of variation in these regions will need to be assessed. SvAnna will benefit from future updates to the HPO resource in this area.

SvAnna runs in about 15-20 minutes for a typical genome on a consumer laptop (faster if computations are performed on more threads). All required data for running SvAnna are provided as a compressed archive for download. SvAnna is implemented as a standalone application with no external dependencies. SvAnna is currently the only tool for phenotype-based prioritization of SVs that is specifically designed to work with VCF files produced by typical LRS SV callers. We are likely to be at the beginning of a period of rapid expansion of LRS in diagnostic settings. SvAnna will play an important role in this process by promoting improved clinical interpretation of a range of SVs. The interpretable prioritizations provided by SvAnna will facilitate the widespread adoption of LRS in diagnostic genomics.

## Methods

### Data sources

The input variants are filtered to remove the common SVs before the prioritization SvAnna uses several sources of common SVs and their frequencies: Database of Genomic Variants,^37^ GnomAD SV,^38^ Human Genome Structural Variation Consortium (HGSVC) SVs freeze 3,^6^ and dbSNP v151 databases (all accessed in April 2021). Transcript definitions were generated for UCSC, RefSeq and Ensembl using Jannovar.^39^ Enhancer definitions were extracted from the VISTA database.^27^ Topologically associating domain (TADs) boundary definitions were taken from a set of evolutionarily conserved TADs that are stable in multiple cell types.^23^ Locations of repetitive elements were taken from the UCSC Genome Database.^26^ Computational disease definitions were extracted from the Human Phenotype Ontology^16^ (HPO) resource (06/2021 release).

### Comprehensive and harmonized representation of variants in VCF files

SvAnna was designed to capture all classes of structural variation represented in VCF files. It transforms default (sequence) and symbolic representations of variants including BND calls into a standardizing computational model of location and consequences. For this purpose, we developed a Java library for representing genomic variants and regions that provides a standardized approach to representing genomic coordinate systems, VCF trimming strategies, and represents small, structural and breakend variants with a consistent API (https://github.com/exomiser/svart).

### Variant prioritization

To prioritize a variant, SvAnna first determines the evaluation region by looking up- and downstream of the variant to find the nearest TAD boundaries. SvAnna uses a threshold of 0.9 for TAD tissue stability^15^ to create a TAD region that is stable across multiple cell types (this parameter can be adjusted if desired). It then analyzes each genomic element contained in the TAD separately. If no boundary is located between an SV and start or end of the chromosome, the start or end position is taken. SvAnna applies different rules for the various categories of SV. For each gene in the TAD, a functional score is calculated to reflect the predicted difference in functionality and gene dosage between the reference and alternate sequence, *δ*(*g*) = |*δ*_*ref*_(*g*) - *δ*_alt_(*g*)|. The rules for calculating *δ*differ according to the SV type and are described in the following sections.

### Deletions, duplications, and inversions

SvAnna evaluates impact of a SV on all genes *g* ∈ *G* present in the TAD region. Scoring *δ(G)* for deletions and duplications depends on the type of sequence affected. The maximal deleteriousness is scored for coding sequences. A deletion that disrupts the coding sequence of a transcript, either because it removes the entire transcript or because it removes part of the coding sequence by deletion of one or more exons, receives a *δ(G)* score of 0. For UTRs, the score *δ*(*g*) for gene *g* is determined as a function of SV and UTR length, with the difference between wildtype (1) and variant sequence calculated as follows:

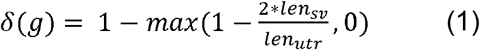

A deletion that encompasses 50% or more of the UTR will be assigned a score of 0 (maximal deleteriousness). Smaller deletions will receive proportionally less deleterious scores. Analogously, a duplication that adds an entire copy of a transcript without disrupting the coding sequence is assigned a *δ(g)* score of 2 (a triplication would be assigned a score of 3, and so on).

If a breakpoint of an inversion disrupts the coding sequence of a transcript, it is assigned a score of 0. If a transcript is completely contained within an inversion, its score is unchanged (1).

### Insertions

To score insertions, SvAnna considers several special cases. For an insertion located within the coding sequence of a transcript, SvAnna checks if the insertion disrupts the reading frame. The insertion that fits into the reading frame is assigned a *δ(g)* score of 0.8 if the number of inserted bases is a multiple of three, or 0.5 if the number of bases is not a multiple of three. Out of frame insertions within the coding sequence are assigned a *δ(g)* score of 0.1. If the insertion is located in UTR, the *δ(g)* score is determined as a function of insertion and UTR length:

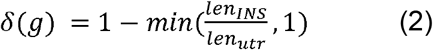

An insertion that adds the number of bases corresponding to 100% of the UTR will be assigned a score of 0 (maximal deleteriousness). The shorter insertions will receive proportionally less deleterious scores. An insertion located outside of the coding sequence, splice site regions, promoter, and UTRs is assigned a score of 1 (no deleteriousness).

### Translocations

If a breakpoint of a translocation disrupts the coding sequence of a transcript, it is assigned a score of 0 (1). The scoring process applies to both breakends of a translocation.

### Phenotype matching

SvAnna takes a list of HPO terms describing the clinical manifestations of the proband as input. It matches them to the 7,981 computational disease models of the HPO using symmetric Resnik matching.^40^ Briefly, to calculate the Resnik symmetric matching score □*(Q, D)* for disease *D* annotated with {*d_1_,…,d_m_*} HPO terms and query *Q* consisting of {*q1,…, q_n_*} HPO terms describing the clinical manifestations of the proband, SvAnna uses:

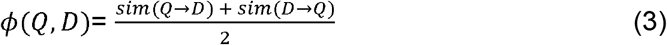

where *sim*(*Q* → *D*) is a method for obtaining one-sided semantic similarity between *Q* and *D.* The one-sided semantic similarity is obtained as:

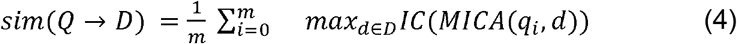

To save computational time, SvAnna pre-calculates the information content of the most informative common ancestor *IC(MICA(t_1_, t_2_))* for all terms *t* used to annotate computational disease models.

### The TAD_SV_ score

The TADwise Assessment of Deleteriousness of Structural variation (TAD_SV_) score is calculated based on the sequence scores *δ(G)* and phenotype similarity scores □*(Q, D)* for all genes *G* contained within the TAD that contains the SV. The sequence score *δ(G),* which represents alterations between wildtype and variant sequences for all genes affected by the SV, is weighted by the phenotypic similarity score □*(Q, D).*

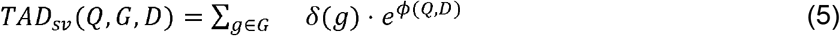

Here, the TAD_sv_ score is calculated as a function of the query HPO terms (*Q*), the genes *G* and the Mendelian diseases *D* associated with the genes in *G.* The function *δ(g)*calculates the difference between wildtype and variant sequences for the variant of interest. *δ(g)* is then weighted by the exponentiated phenotypic similarity *□(Q, D)* of the query terms *Q* to a computational model of a disease *D* that is associated with variants in *g.* SvAnna uses the highest □*(Q, D)* if more than one disease is associated with variants in *g*.

### Variant filtering

In our analyses, we considered the variants occurring in more than 1% of the population as common. SVs called in the VCF file are removed from the analysis if they show greater than 80% reciprocal overlap with a common variant in any of the source databases (DGV, gnomAD-SV, dbSNP, and HGSVC SVs). The frequency and reciprocal overlap thresholds may be adjusted via the command line interface.

### SvAnna performance benchmarks and comparison of with other algorithms for ranking pathogenic SVs

We compared SvAnna with AnnotSV.^23^ Since all the tools evaluate each variant independently, we used a simple strategy to simulate a variant dataset, by adding in variants from published cases of disease-associated SVs (Table 1) to VCF files derived from PacBio whole-genome sequencing and called with pbsv (see below). Each published variant was added in turn to one of ten in-house VCF files produced in this way. Each of the two algorithms was applied to the cases and the median rank from the ten VCF files was recorded. Then the benchmarking study was performed using the pipeline engine Nextflow.^41^

### AnnotSV

*AnnotSV* is an open source tool that annotates structural variants stored in VCF and BED format with functional, regulatory, and clinical information and classifies SVs into five pathogenicity classes: *benign* (1), *likely benign* (2), *variant of uncertain significance* (3), *likely pathogenic* (4) and *pathogenic* (5). *AnnotSV* reports results in the tabular format. We performed a local installation of *AnnotSV* v3.0.9 (accessed on June 25 2021). We used *full* annotation mode when running *AnnotSV* to produce a single row per SV in the result file, and we set the *-SVminSize* to 1 to ensure SVs shorter than 50bp (including causal SVs) are analyzed. The tabular output reports the annotated SVs as one SV per row, and the rows are ordered by decreasing priority. We used the row number to determine the variant rank. If the causal variant was assigned *pathogenicity class=NA* instead of one of the five pathogenicity classes, and therefore missed, we assigned the variant the rank 40,000.

### Long read sequencing

We used VCF files from ten in-house whole genome sequencing experiments as background files for the simulations. We do not have consent for sharing the files.

#### High molecular weight (HMW) DNA extraction

The HMW gDNA was extracted using the Gentra Puregene (Qiagen) kit. Frozen cells or tissues were first pulverized using a mortar and pestle and transferred to a 15mL tube that contained Qiagen Cell Lysis Solution. The lysate was then incubated with Proteinase K for 3 hours at 55°C, followed by RNase A for another 40 minutes at 37°C. Samples were cooled on ice and Protein Precipitation Solution was added. Samples were then vortexed and centrifuged. The supernatant was transferred to a new tube containing isopropanol for precipitation. Pellet was washed with 70% ethanol, air dried, rehydrated in PacBio Elution Buffer until dissolved.

#### PacBio HiFi whole genome sequencing

This protocol was carried out using the PacBio SMRTbell Express Template Prep Kit 2.0 and the SMRTbell Enzyme Cleanup Kit. 15 μg of DNA was sheared to 20kb using g-TUBE (Covaris). The sheared DNA was purified using Ampure PB beads (PacBio). 10 μg of sheared DNA was used in removing single strand overhangs, followed by DNA damage repair and End repair/ A-tailing. The repaired/ modified DNA was used for V3 adapter Ligation. The adapter ligated library was treated with Enzyme mix for Nuclease treatment to remove damaged or non-intact SMRTbell templates. The purified library was then size selected using two-step size selection with Blue Pippin (Sage Science) generating 9-13kb and >15kb fractions. The size selected and purified >15kb fraction of the library was used for sequencing on Sequel II.

#### Alignment and variant calling

We aligned long-read PacBio HiFi WGS data to the GRCh38 (hg38) reference using pbmm2 (v1.3.0) with *--preset CCS* option enabled. We identified SVs on the indexed bam files using pbsv(v2.4.0).

### URLs

- SvAnna source code - https://github.com/TheJacksonLaboratory/SvAnna
- SvAnna tutorial and documentation - https://svanna.readthedocs.io
- Global Alliance for Genomics and Health (GA4GH) Phenopackets (v1) representing the published case reports of deleterious SVs - *https://zenodo.org/record/5071267*
- Database of Genomic variants (DGV) - http://dgv.tcag.ca
- GnomAD SV - https://ftp.ncbi.nlm.nih.gov/pub/dbVar/data/Homo_sapiens/by_study/genotype/nstd166/gnomad_v2.1_sv.sites.vcf.gz
- HGSVC SV - http://ftp.1000genomes.ebi.ac.uk/vol1/ftp/data_collections/HGSVC2/release/v1.0/integrated_callset/freeze3.sv.alt.vcf.gz
- dbSNP - https://ftp.ncbi.nih.gov/snp/organisms/human_9606_b151_GRCh38p7/VCF/00-common_all.vcf.gz
- TAD boundary definitions - https://zenodo.org/record/4156731/files/emcarthur/TAD-stability-heritability-v1.0.1.zip
- PacBio structural variant (SV) calling and analysis tools. https://github.com/PacificBiosciences/pbsv
- PacBio pbmm2 aligner: https://github.com/PacificBiosciences/pbmm2

## Ethics declarations

Competing interests: The authors declare no competing interests.

## Code availability

SvAnna code is freely available for academic use on GitHub (see URLs).

## Data availability

SvAnna database files are available for download via the SvAnna documentation site (see URLs). The case reports of deleterious SVs are available for download from Zenodo (see URLs).

