## Supplemental Figures for "SvAnna: efficient and accurate pathogenicity prediction for coding and regulatory structural variants in long-read genome sequencing"

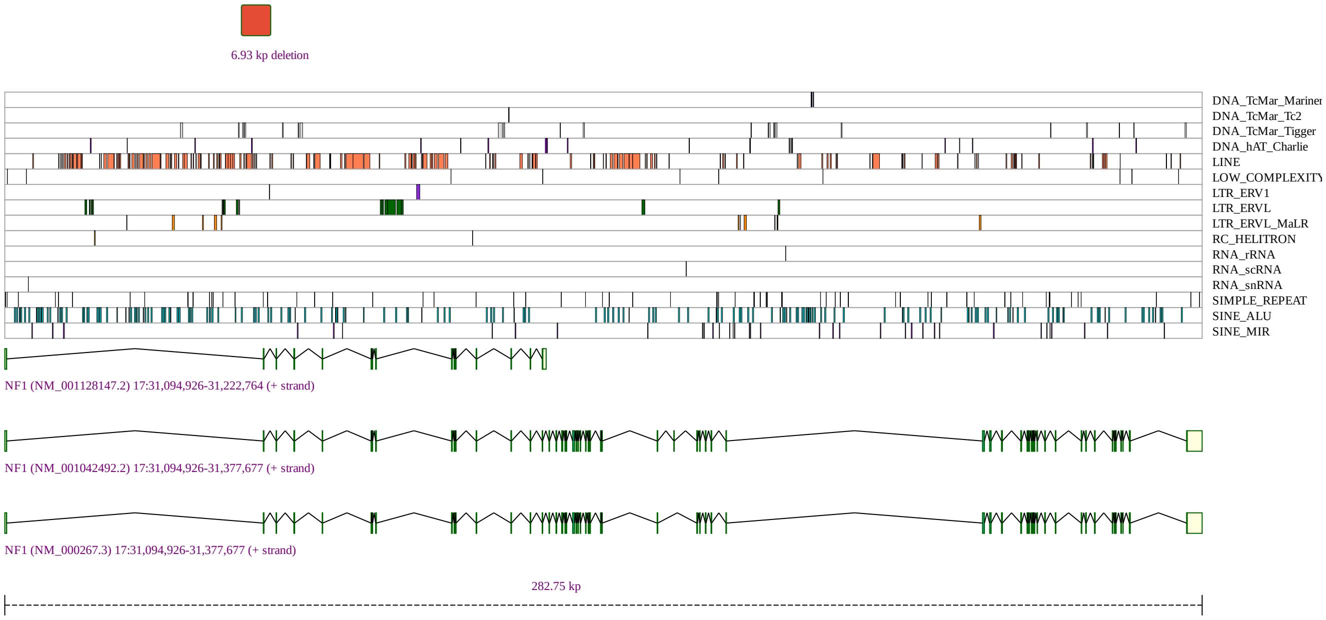


**Figure S1. Deletion affecting** *NF1*. Screenshot of the graphic generated by SvAnna for chr17:31,150,798-31,157,725del, a deletion of 6.93 kb that was assigned a priority score of 124.63. The deletion affects exon 2 of several *NF1* transcripts including NM_000267.3. Pathogenic variants in *NF1* are associated with neurofibromatosis type 1 (OMIM:162200). The phenotypic features curated for this case were *Multiple cafe-au-lait spots* (HP:0007565), *Plexiform neurofibroma* (HP:0009732), *Spinal neurofibromas* (HP:0009735), and *Tibial pseudarthrosis* (HP:0009736). Data were curated from a published case report [1].

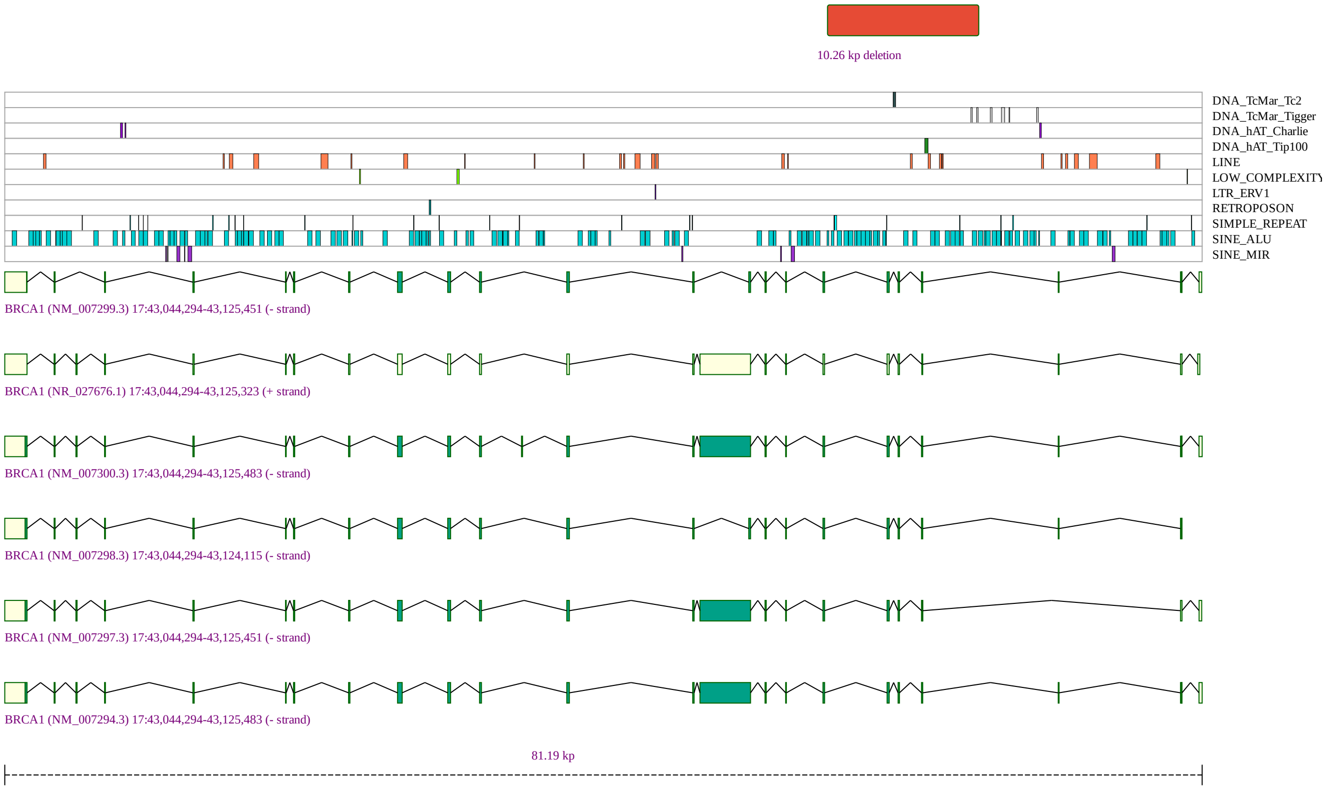


**Figure S2. Deletion affecting** *BRCA1*. Screenshot of the graphic generated by SvAnna for chr17:43,100,079-43,110,335del, a deletion of 10.26 kb that was assigned a priority score of 66.41. The deletion affects three *BRCA1* exons. Pathogenic variants in *NF1* are associated with Breast-ovarian cancer, familial, 1 (OMIM:604370). The phenotypic feature curated for this case was *Breast carcinoma* (HP:0003002). Data were curated from a published case report [2].

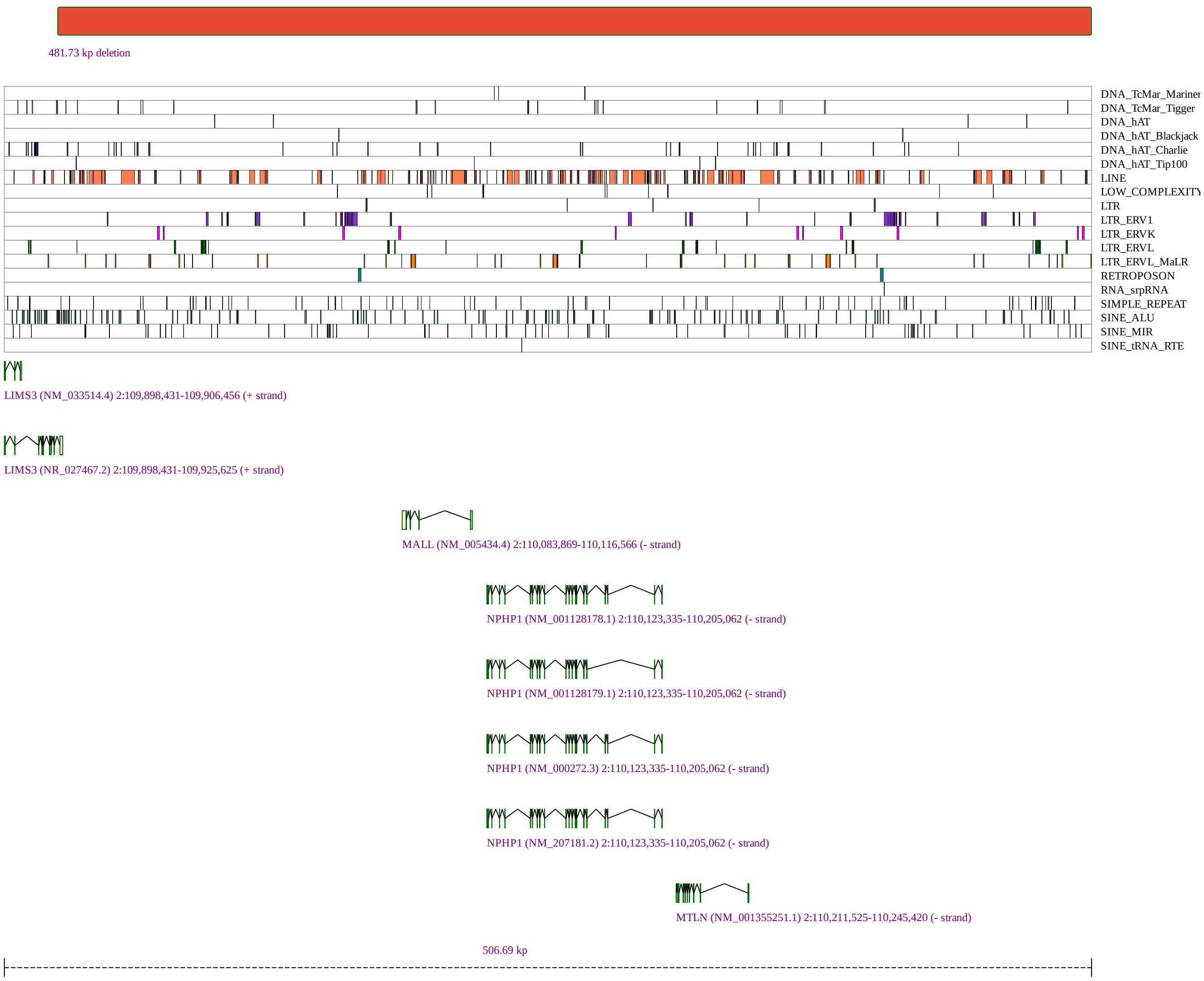


**Figure S3. Deletion affecting** *NPHP1*. Screenshot of the graphic generated by SvAnna for chr2:109,923,337-110,405,062del, a deletion of 481.73 kb that was assigned a priority score of 15.14. The deletion is predicted to disrupt *MTLN*, *MALL*, *MTLN*, and *NPHP1*. Pathogenic variants in *NPHP1* are associated with Joubert syndrome 4 (OMIM:609583). The phenotypic features curated for this case were *Stage 5 chronic kidney disease* (HP:0003774), *Cerebellar vermis hypoplasia* (HP:0001320), *Truncal ataxia* (HP:0002078), *Blindness* (HP:0000618), *Ptosis* (HP:0000508), *Molar tooth sign on MRI* (HP:0002419), *Elongated superior cerebellar peduncle* (HP:0011933), *Limb ataxia* (HP:0002070, *Optic disc pallor*(HP:0000543), and *Coloboma* (HP:0000589). Data were curated from a published case report [3].

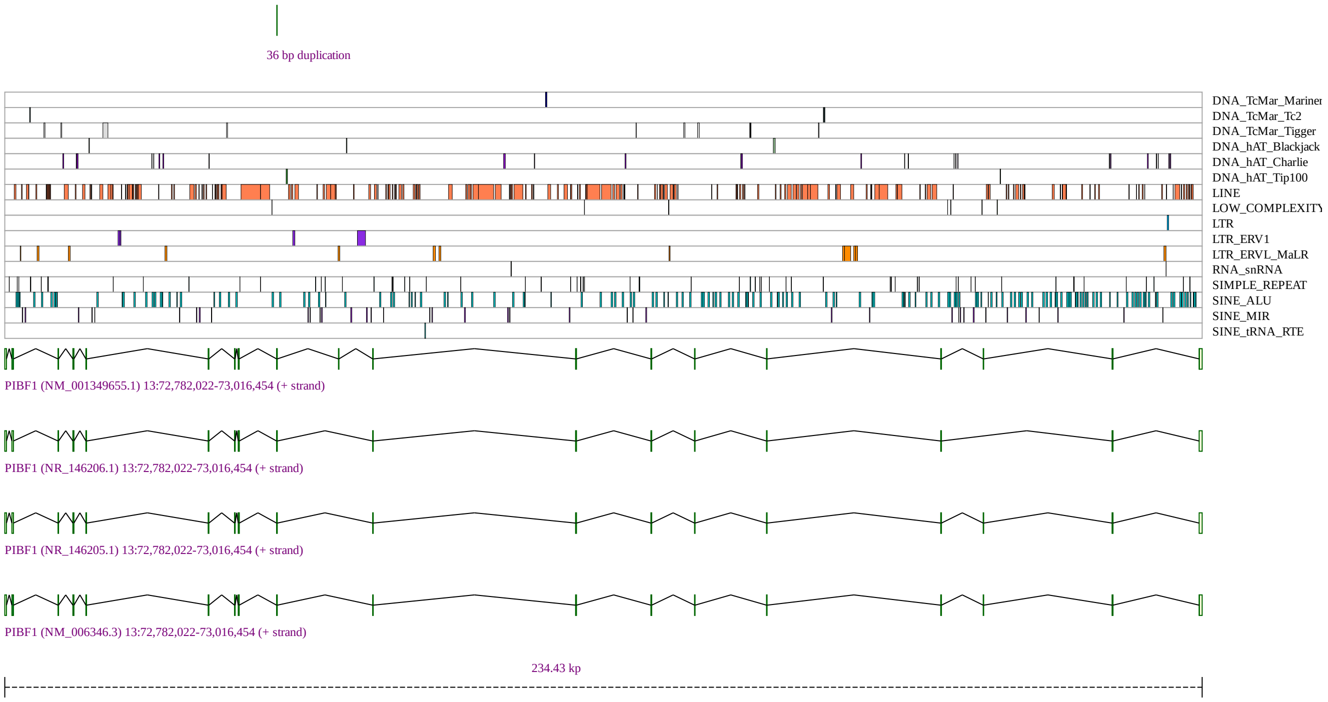


**Figure S4. Duplication affecting** *PIBF1.* Screenshot of the graphic generated by SvAnna for `chr13:72835296-72835332dup`, a duplication of 36 bp that was assigned a priority score of 2.36. Pathogenic variants in PIBF1 are associated with *Joubert syndrome 33* (OMIM:617767). The phenotypic features curated for this case were *Periglomerular fibrosis* (HP:0032417),*Vesicoureteral reflux* (HP:0000076 *Hypoplasia of the corpus callosum* (HP:0002079), *Ascites* (HP:0001541), *Hypermetropia* (HP:0000540), *Feeding difficulties* (HP:0011968), *Seizure* (HP:0001250 *Deeply set eye* (HP:0000490), *Global developmental delay* (HP:0001263), *Areflexia* (HP:0001284), *Hepatomegaly* (HP:0002240), *Generalized hypotonia* (HP:0001290), *Hyaline casts* (HP:0031200), *Midface retrusion* (HP:0011800), *Nephronophthisis* (HP:0000090), *Renal tubular atrophy* (HP:0000092), *Acute kidney injury* (HP:0001919 *Perisylvian polymicrogyria* (HP:0012650), *Molar tooth sign on MRI* (HP:0002419 *Ventriculomegaly* (HP:0002119), and *Enlarged kidney* (HP:0000105

. Data were curated from a published case report [4].

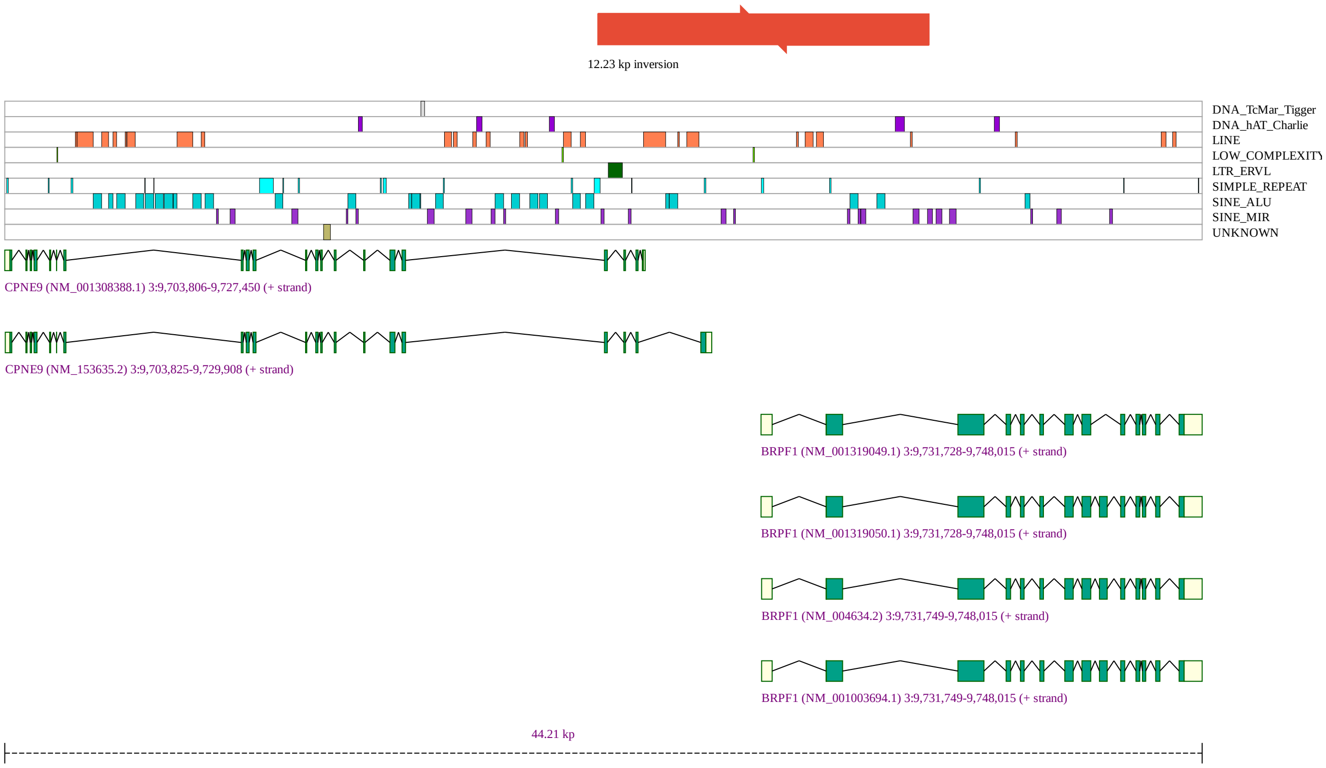


**Figure S5. Inversion affecting** *BRPF1*. Screenshot of the graphic generated by SvAnna for `inv(chr3)(9725702; 9737931)`, a ∼12.23 kb inversion that disrupts the coding sequence of the *BRPF1* gene with a Priority score of 7.76. Pathogenic variants in *BRPF1* are associated with *Intellectual developmental disorder with dysmorphic facies and ptosis* (OMIM:617333. The phenotypic features curated for this case were *Epicanthus* (HP:0000286), *Bilateral tonic-clonic seizure* (HP:0002069), *Downslanted palpebral fissures* (HP:0000494), *Intellectual disability, moderate* (HP:0002342), *Strabismus* (HP:0000486), *Delayed speech and language development* (HP:0000750), *Wide nasal bridge* (HP:0000431), *Hypotonia* (HP:0001252), *Delayed gross motor development* (HP:0002194), *Flat face* (HP:0012368), *Myoclonic absence seizure* (HP:0011150), *Fused cervical vertebrae* (HP:0002949), *Ptosis* (HP:0000508), *Hypertelorism* (HP:0000316), and *Round face* (HP:0000311). Data were curated from a published case report [5].

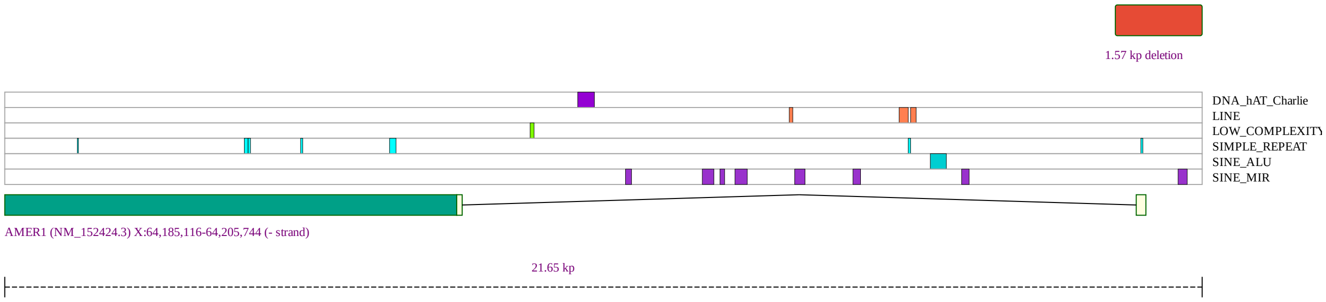


**Figure S6. Deletion affecting** *AMER1*. Screenshot of the graphic generated by SvAnna for chrX:64,205,190-64,206,761del, a hemizygous deletion of ∼1.57 kb that deletes the transcription start site of the *AMER1* gene and was assigned a Priority score of 9.02. Pathogenic variants in *AMER1* are associated with Osteopathia striata with cranial sclerosis (OMIM:300373). The phenotypic features curated for this case were *Polyhydramnios* (HP:0001561), *Delayed speech and language development* (HP:0000750), *Thickened calvaria* (HP:0002684), *Upper airway obstruction* (HP:0002781), *Hypertelorism* (HP:0000316), *Metaphyseal striations* (HP:0031367), *Bilateral cleft lip and palate* (HP:0002744), *Macrocephaly* (HP:0000256), and *Lymphedema* (HP:0001004). Data were curated from a published case report [6].

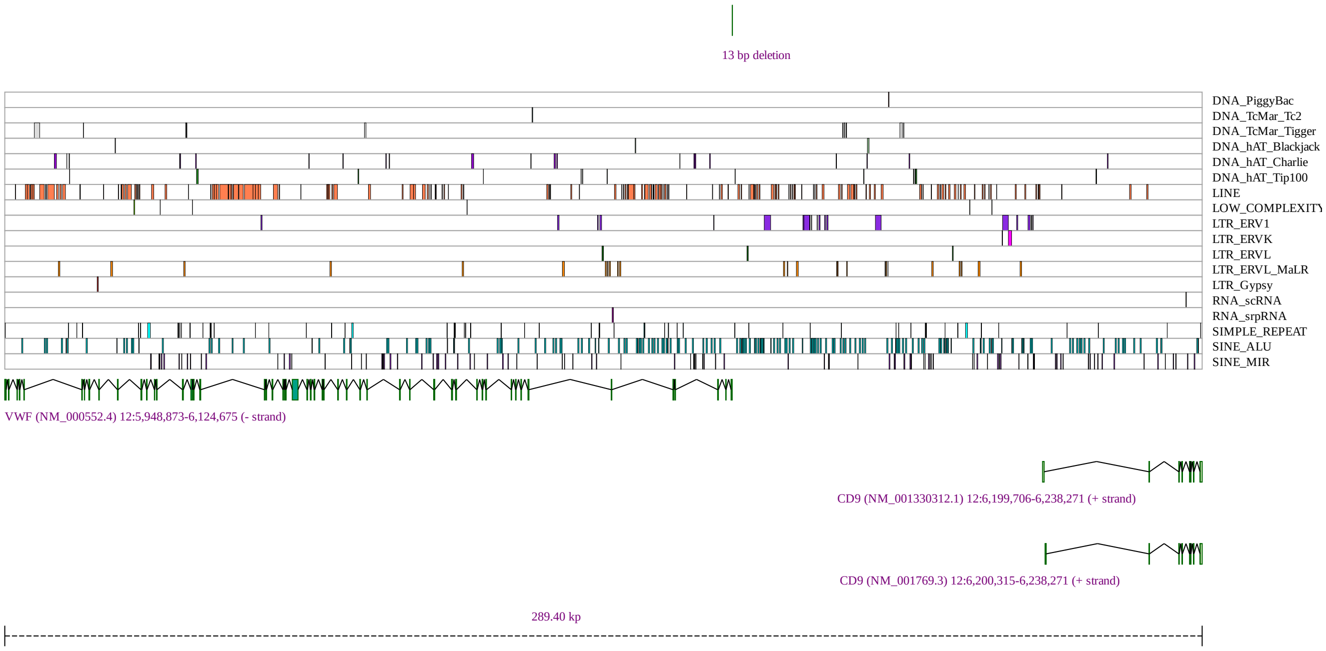


**Figure S7. Deletion affecting** *VWF*. Screenshot of the graphic generated by SvAnna for chr12:6,124,705-6,124,718del, a 13 bp deletion located in the core promoter region of *VWF* ( 30 bp upstream of NM_000552.4). In the original publication, the deletion was shown to lead to aberrant binding of Ets transcription factors to the site of the deletion and thereby reduce VWF expression [7]. Pathogenic variants in *VWF* are associated with von Willebrand disease. The phenotypic features curated for this case were *Prolonged bleeding following procedure* (HP:0011890), *Bruising susceptibility* (HP:0000978), *Reduced quantity of Von Willebrand factor* (HP:0012147). Data were curated from a published case report [7].

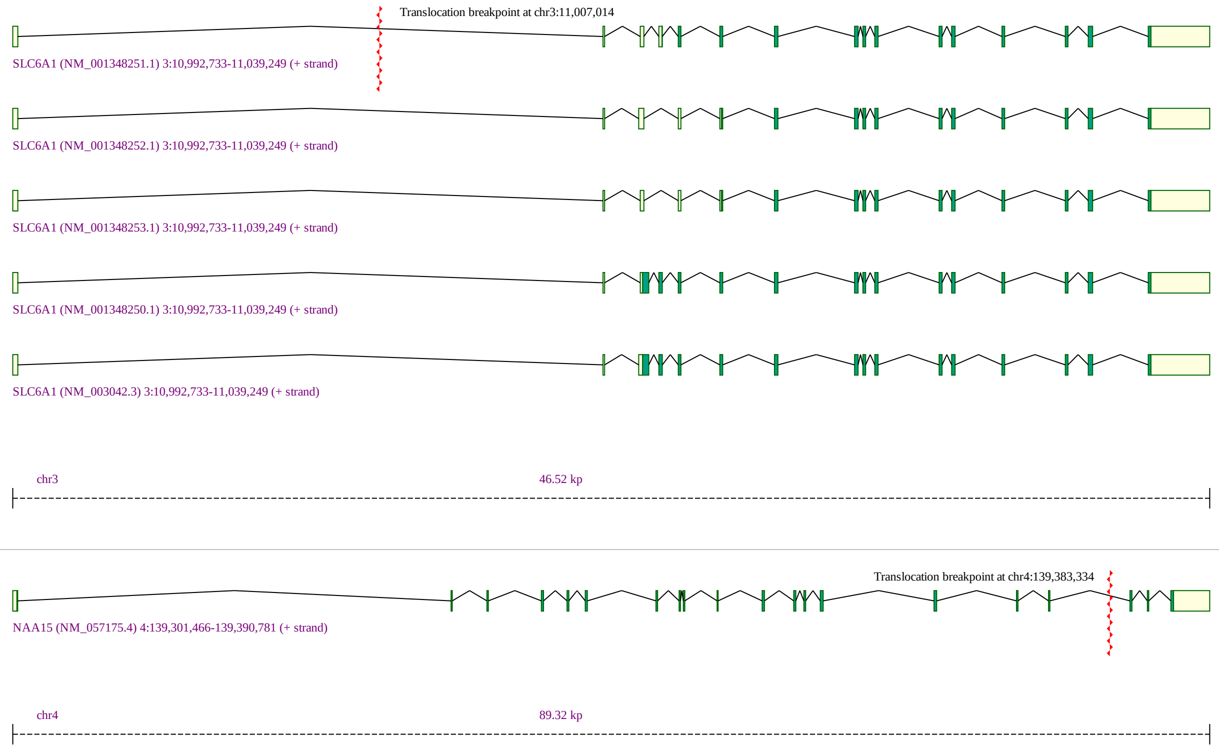


**Figure S8. Translocation affecting** *SLC6A1*. Screenshot of the graphic generated by SvAnna for `t(chr3:11,007,014; chr4:139,383,334)`, a translocation that was assigned a priority score of 4.63. Pathogenic variants in *SLC6A1* are associated with *Myoclonic-atonic epilepsy* (OMIM:616421). The phenotypic features curated for this case were *Hypertonia* (HP:0001276), *Microcephaly* (HP:0000252), *Knee flexion contracture* (HP:0006380), *Scoliosis* (HP:0002650), *Seizure* (HP:0001250), *Global developmental delay* (HP:0001263), *Malar flattening* (HP:0000272), *Thin upper lip vermilion* (HP:0000219), *Thick lower lip vermilion* (HP:0000179), *Elbow flexion contracture* (HP:0002987), *Narrow nasal bridge* (HP:0000446). Data were curated from a published case report [8].

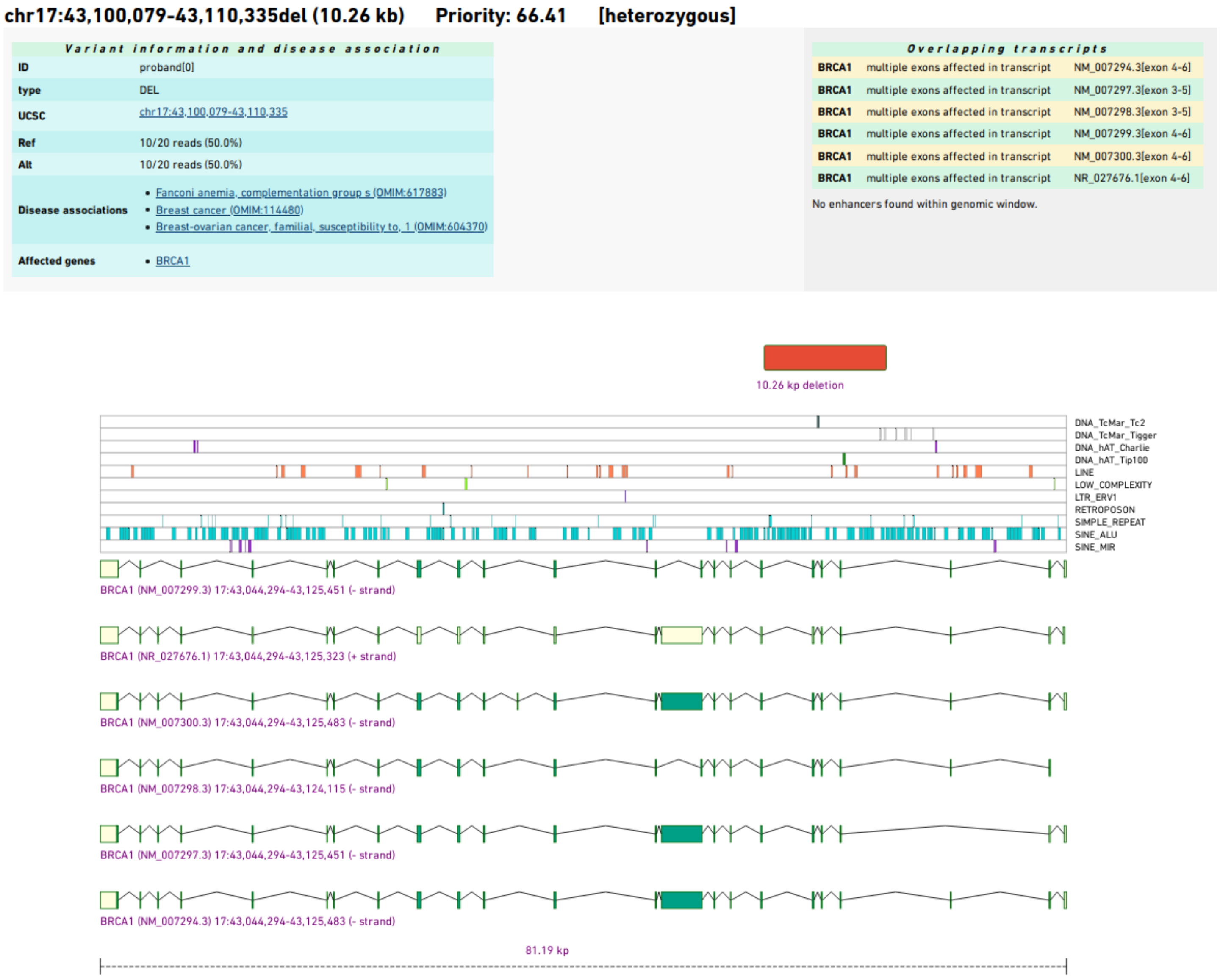


**Figure S9.** SvAnna screenshot. SvAnna generates a tabular and graphical summary of the top 100 variants (the number can be adjusted by the user). The chromosomal position, length, priority score, genotype are listed at the top of each entry. The *Variant information and disease association* table summarizes read count information and the ID of the variant in the VCF file, and provides links to genes affected by the SV and diseases associated with the gene in the Online Mendelian Inheritance in Man (OMIM) resource [9]. The *Overlapping transcripts* table shows each of the transcripts that overlaps with the SV as well as the position and effect on the transcript. The graphical display is generated as a scalar vector graphics (SVG) file that is embedded directly in the HTML code. It shows the SV and its position compared to that of overlapping transcripts, whose coding exons are shown in green and non-coding exons in yellow. If applicable, overlapping repeats are shown as tracks beneath the variant. In some cases, this may provide a clue as to whether a given SV event could be related to neighboring repetitive sequences. Information for this track is derived from the UCSC Genome Browser database [10].

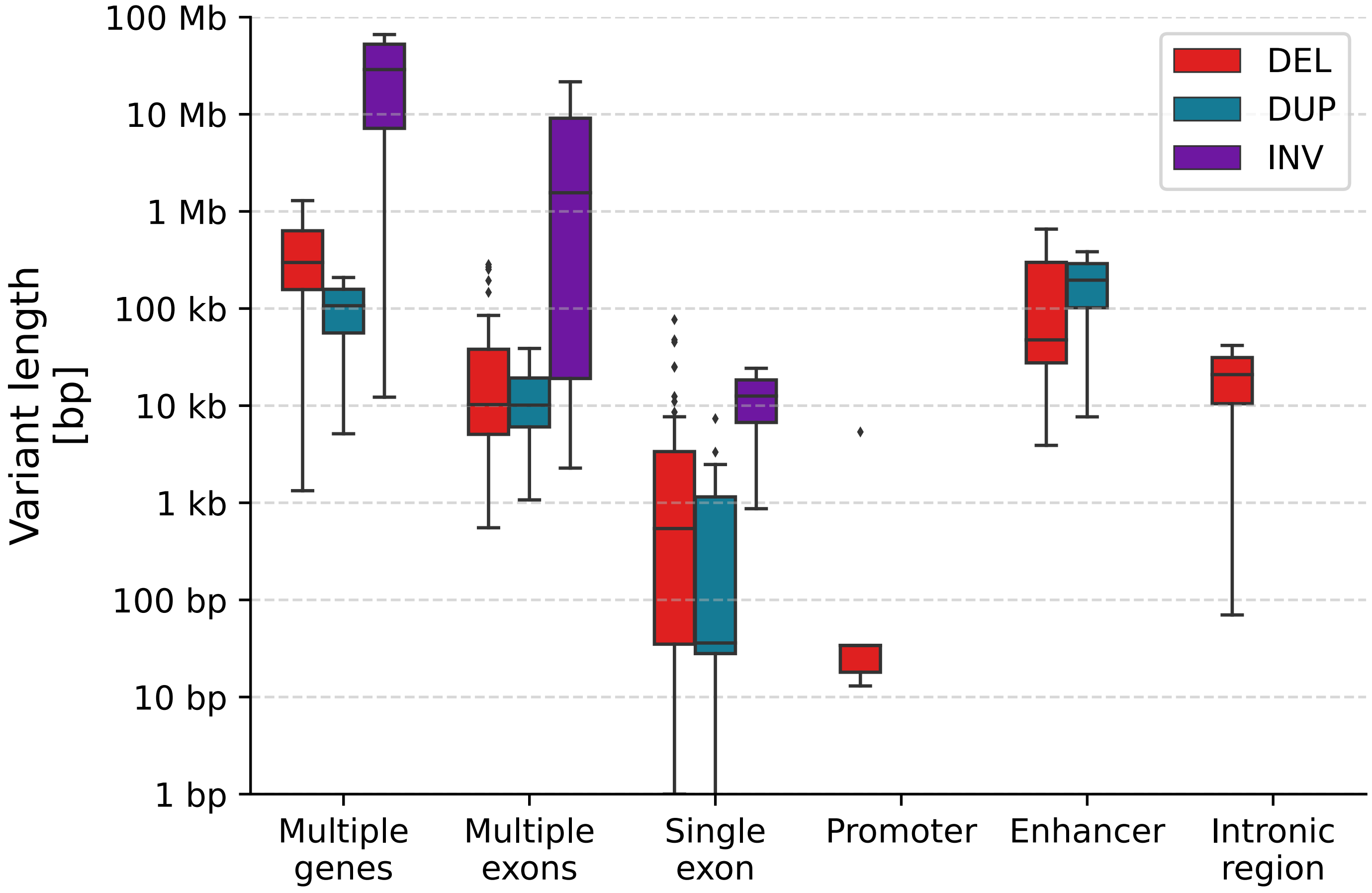


**Figure S10. Distribution of lengths of the curated structural variants**. Insertions and translocations are considered to have zero length (not shown).
